# Functional consequences of shifting transcript boundaries in glucose starvation

**DOI:** 10.1101/2023.07.02.547342

**Authors:** Lan Anh Catherine Nguyen, Masaru Mori, Yuji Yasuda, Josephine Galipon

## Abstract

Glucose is a major source of carbon and essential for the survival of many organisms, ranging from yeast to human. A sudden 60-fold reduction of glucose in exponentially growing fission yeast induces transcriptome-wide changes in gene expression. This regulation is multilayered, and the boundaries of transcripts are known to vary, with functional consequences at the protein level. By combining direct RNA sequencing with 5’-CAGE and short-read sequencing, we accurately defined the 5’- and 3’-ends of transcripts that are both poly(A) tailed and 5’-capped in glucose starvation, followed by proteome analysis. Our results confirm previously experimentally validated loci with alternative isoforms and reveal several transcriptome-wide patterns. First, we show that sense-antisense gene pairs are more strongly anti-correlated when a time lag is taken into account. Secondly, we show that the glucose starvation response initially elicits a shortening of 3’-UTRs and poly(A) tails, followed by a shortening of the 5’-UTRs at later time points. These result in domain gains and losses in proteins involved in the stress response. Finally, the relatively poor overlap both between differentially expressed genes (DEGs), differential transcript usage events (DTUs), and differentially detected proteins (DDPs) highlight the need for further study on post-transcriptional regulation mechanisms in glucose starvation.

## Introduction

Glucose is a major source of carbon used by many organisms to generate energy for cell multiplication and survival. Nevertheless, previous studies have linked an excess of glucose with metabolic disorders such as inflammation or diabetes that might induce shorter life-span and promote the multiplication of tumoral cells.^1, 2^ In contrast, glucose restriction has been associated with several health advantages including an increased life span in most model organisms including fission and budding yeast,^3, 4^ worm,^5^ mouse and fly.^6, 7^ Most recently, direct RNA sequencing (DRS) in *Schizosaccharomyces pombe* revealed alternative splicing events,^8^ confirming the fission yeast as a model of interest for the study of post-transcriptional regulation.^9^ Transcriptional shifts in stress responses such as osmotic stress, oxidative stress, and heat shock have been studied at the transcriptional level,^10^ but the response to glucose starvation remains relatively understudied. When exposed to glucose-deprived conditions, fission yeast switches from the glucose-consuming glycolysis pathway to the glucose-producing gluconeogenesis pathway.^11^ In particular, while previous studies have investigated gene expression patterns in fission yeast exposed to glucose starvation,^12–15^ the extent to which the boundaries of transcripts shift during the response is poorly understood. The 5’- and 3’-ends of transcripts define the length and therefore the sequence composition of 5’- and 3’-Untranslated Regions (UTRs) with implications on RNA stability and coding potential.^16, 17^

This study investigates the changes in transcriptional start and end sites in the fission yeast *Schizosaccharomyces pombe* 972h-during glucose starvation to get better mechanistic insight on the transcriptional dynamics. When exposed to glucose starvation stress, the fission yeast undergoes a stress response that manifests itself at several levels of gene regulation. Previous studies have shown that TORC2 (target of rapamycin complex 2),^18^ and cAMP (cyclic AMP) signaling pathways are activated under limited glucose availability. While the former is essential for cell proliferation under glucose-limited conditions, the latter activates the protein kinase A (PKA) for inhibiting glucose starvation-induced processes, including the transcription of the *fbp1* gene that codes for fructose-1,6-biphosphatase, one of the key enzymes in the gluconeogenesis pathway.^19^ Upon glucose starvation, same-strand transcripts overlapping the entire *fbp1*+ locus are transcribed starting from the region upstream from the promoter, but do not produce any protein.^12^ These transcripts were subsequently defined as metabolic long non-coding RNA (mlonRNA)^13^(p201) and were shown to participate in the full activation of *fbp1* mRNA transcription by directly recruiting transcription factors.^20^ These transcripts, as well as their antisense, harbor both poly(A) tail and 5’-cap. They are also bound by multiple ribosomes in the cytoplasm despite being non-coding, suggesting a role of RNA surveillance pathways in RNA selection during stress.^13, 15^ Some of these transcripts may be seen as isoforms with an alternative transcription start site (aTSS), and may be examples of what is now known as differential transcript usage (DTU).^21^ Short-read sequencing technology has a hard time identifying these events with precision, as it is based on RNA fragmentation.^14^

Moreover, most of the data in the database of reference for the fission yeast, Pombase,^22^ still relies on short-read sequencing. Therefore, these alternative transcripts, although partially described in the literature, have yet to be referenced. With long-read sequencing, such as DRS, poly(A) tailed transcripts may be sequenced from end to end, partly overcoming this problem. The principle of this method is the native RNA going through a protein nanopore with an applied voltage. Nucleotides passing sequentially through the pore induce changes in the ionic current. Using a neural network trained on known sequences, it is possible, in theory, to precisely define the sequence of the RNA from the 3’-end to the 5’-end including the poly(A) tail length. This is important as 5’- and 3’-UTRs embed multiple regulatory units such as secondary structures,^23, 24^ enriched motifs, RNA binding proteins binding sites, and small Open Reading Frames (ORFs) in human, called upstream ORFs (uORFs) for 5’-UTRs and downstream ORFs for 3’-UTR (dORFs), and potentially code for small peptides with functional sequences.^25–27^ In other words, there is a need to use new technologies to precisely define the 5’- and 3’-boundaries of transcripts expressed by the fission yeast, and especially during glucose starvation to better understand how these noncoding regions participate in gene regulation.

Although DRS is suitable for identifying transcript isoforms along with the position of transcription termination and the length of the poly(A) tail, the accuracy of basecalling remains the lowest among existing methods. In addition, precise mapping of the 5’-ends of transcripts is relatively more challenging due to premature runoff of some transcripts prior to reaching the 5’-end.^28^ In addition, the last 10 to 15 nucleotides at the 5’-end cannot be read.^29^ To work around these limitations, we sequenced the whole transcriptome with shotgun cDNA sequencing (formerly known as RNA-Seq) and the 5’-ends of transcripts using Cap Analysis of Gene Expression sequencing (CAGE-sequencing).^30^ By leveraging the combined advantages of the above three sequencing methods, we reannotated the fission yeast transcriptome in response to glucose starvation with unprecedented accuracy. This revealed two poorly overlapping sets of differentially regulated transcripts: DEGs, and transcripts with shifted boundaries or DTUs. Considering DEG-antisense pairs, 25.1% and 31.9% were positively and negatively correlated with their antisense expression, respectively. In addition, we show 3’-UTRs and poly(A) tails shorten early in glucose starvation, followed by a shortening of the 5’-UTRs with functional consequences on protein domains. All in all, this study constitutes the first example of mapping the precise 5’- and 3’-boundaries of transcripts that are both capped and polyadenylated in glucose starvation.

## Material and Methods

### Strains pre-culture and culture

20 mL of cells were harvested for time point 0 minutes from the Yeast Extract Rich (YER) medium (0.5% w/v yeast extract; 6% glucose; 100 mg/L adenine) when the concentration reached 1.8–2.2 × 10^7^ cells/mL. The remaining cells were transferred to Yeast Extract Dextrose (YED) medium (0.5% w/v yeast extract; 0.1% glucose; 3% glycerol; 100 mg/L adenine) after washing them twice with distilled water. 20 mL of the cell cultures were collected by centrifugation at different time points (15 minutes, 60 minutes and 120 minutes), and rinsed once in ice-cold 1X Phosphate Buffered Saline (PBS) (137 mM NaCl; 2.7 mM KCl; 10 mM Na2HPO4; 1.8 mM KH2PO4). After sampling, the cells were flash-frozen in liquid nitrogen and stored at −80°C.

### RNA extraction

Frozen pellets were re-suspended in 250 µL of 65°C sterilized bead buffer (7.5 mM NH4OAc; 10 mM EDTA, pH 8.0) and then transferred to 1.5 mL tubes prewarmed at 65°C acid-washed glass beads (G8772-100 Merck) suspended in 25 µL of 10% filtered SDS and 300 µL of acid phenol chloroform pH 4.5. The tubes were vortexed for 1 minute then incubated at 65°C for 1 minute. This was repeated three times. After heating the tubes at 65°C for 10 minutes, and after an additional 1 minute-round of vortex, the samples were centrifuged at 16,000 x g, 25°C, for 15 minutes. The supernatant was then transferred to a new tube with an equal volume of chloroform on ice. After vortexing the tubes, the tubes were centrifuged at 16,000 x g, 4°C, for 15 minutes. The supernatant was transferred to a new tube with 7.5 M NH4OAc (final concentration of 0.51M). The tubes were then vortexed. After adding 100% cold ethanol, the tubes were put at −80°C. The samples were then centrifuged at 20,400 x g at 4°C for 30 minutes. After discarding the supernatant, the pellet was washed in 75% ice-cold ethanol was then added. After a final centrifugation at 20,400 x g, 4°C, for 15 minutes, the supernatant was discarded. The pellet was dried using a vacuum-centrifuge for 5 minutes and re-suspended on ice with cold RNAse-free water and stored at −80°C. RNA quality and concentration was measured using a Nanodrop spectrophotometer (ND-1000).

### Direct RNA library preparation and sequencing

RNA library preparation was done using direct RNA sequencing (DRS) kit (SQK-RNA002 from Oxford Nanopore Technology following the manufacturer’s instructions. The protocol was slightly modified: SuperScript IV reverse transcriptase (Invitrogen) was used instead of SuperScript III. The samples were incubated in a thermocycler at 50°C for 35 minutes instead of 50 minutes, followed by inactivation at 70°C for 15 minutes instead of 10 minutes before bringing the samples to 4°C. DRS was then done on the flowcell FLO-MIN106 using Oxford Nanopore Technology (ONT) MinION sequencer according to the manufacturer’s protocol.

### Short-read sequencing using BGI technology

Total RNA extracted following the protocol described in the previous section was outsourced to BGI Japan. Total RNA quality was assessed by Bioanalyzer (Agilent) following the manufacturer’s protocol. After mRNA enrichment by oligo dT selection, the mRNA was fragmented and reverse transcribed. The second strand was synthesized, and the ends were repaired with A-tailing, resulting in double-stranded cDNA. Bubble adapters were then ligated. Fragments were amplified using PCR, heat-denatured into single strands and circularized. DNA nanoballs were synthesized and sequenced on a DNBSEQ-G400 platform.

### Cap Analysis of Gene Expression sequencing (CAGE-seq)

Transcription start sites were defined using CAGE sequencing. CAGE library preparation, sequencing, mapping and quality filtering were outsourced to DNAFORM (Yokohama, Kanagawa, Japan). Total RNA extracted following protocol described in 2.3 was sent to DNAFORM. Total RNA quality was assessed by Bioanalyzer (Agilent) following the maker’s protocol. cDNAs were synthesized from total RNA using random primers. Ribose diols in the 5’ cap structures of RNA were oxidized, then biotinylated. The biotinylated RNA/cDNAs were selected by streptavidin beads (cap-trapping). After RNA digestion by RNaseONE/H and adaptor ligation to both ends of cDNA, double-stranded cDNA libraries (CAGE libraries) were constructed.

CAGE libraries were sequenced using single end reads of 75 nucleotides on a NextSeq 500 instrument (Illumina). The obtained reads (CAGE tags) were mapped to the genome of *Schizosaccharomyces pombe* 972h-from Pombase using BWA (v0.5.9).^31^ Unmapped reads were then mapped by HISAT2 (v2.0.5).^32^

### Transcriptome reannotation

Long reads from Nanopore sequencing was basecalled with ONT Guppy GPU Basecaller (v5.0.7). The quality of the reads were controlled with pycoQC (v2.5.2).^33^ The reads were aligned to the reference genome of *Schizosaccharomyces pombe* 972h-using minimap2 (v2.17).^34^ Reads were then converted to SAM and BAM files with the samtools suite (v1.14).^35^

Raw reads quality for short reads was assessed using FastQC v0.11.9. The reads were then aligned to the genome of *Schizosaccharomyces pombe* 972h-using STAR aligner (v2.7.9).^36^

Mapped CAGE reads obtained from the company were converted from bam to bed using the bedtools suite (v2.30.0).^37^ They were then analyzed using the CAGEfightR Bioconductor package (v1.14.0).^38^

The transcriptome was reannotated using FLAIR (v1.5),^39^ with a combination of the long (Nanopore) and short reads (BGI) described above. The annotation obtained by FLAIR was then corrected using SQANTI3 (v4.2),^40^ by providing CAGE tags to validate the position of the 5’ ends. Gene and transcripts were then renamed using IsoAnnot Lite from tappAS pipeline (v1.0.7).^41^ Using the default filter, a transcript is described as an artifact if ‘in the 20bp downstream the annotated Transcription Terminating Site (TTS) there are 12 or more adenines at the genomic level’, ‘3’ end is an intrapriming artifact’, ‘has a junction that is labeled as RT-Switching’, ‘all junctions are either non canonical or does not have a short read coverage above [the given] threshold’ (citation from the SQANTI user manual).^40, 42^

### Transcriptome analysis

Coverage depth and breadth for short reads was assessed using salmon (v1.6.0) with the new transcriptome.^43^ For long reads, it was estimated using NanoCount (v1.1.0).^44^ Counts from NanoCounts were inputted into the DESeq2 package (v1.34.0) for differential gene expression analysis.^45^ Heatmaps were obtained using the ggplot2 package (v3.3.5) for R.^46^ Gene ontology enrichment analysis was performed using clusterProfiler (v3.10.1),^47^ enrichplot (v1.14.1), DOSE (v2.4.0) and ShinyGO (v0.77).^48, 49^

Alternative transcript isoforms were NanoCount quantification files as input. Firstly, isoform switches are identified and ORFs are analyzed. From the resulting amino acid sequences, protein domains, coding potential, signal peptides and disordered regions are determined with CPC2,^50^(p2) IUPred2A,^51^ hmmscan,^52^ and SignalP.^53^ Poly(A) tail length were estimated using nanopolish (v0.14.0).^29^ Wilcoxon signed-rank test was calculated to compare poly(A) tail lengths with scipy package (v1.10.1) with the alternative hypothesis being ‘less’.^54^

### Calculation of the correlation score between sense and antisense gene expression

The compositional variance is described as a ‘cumulative measure of how far the number in a time series spread from the averages of the part they belong in a composition’, while the compositional covariance is a ‘measure of how changes in the part averages of a time series are associated with changes in the part averages of a second time series’.^55^(p32) Combining the equations for both, the compositional correlation score rc is defined by Equation (1):

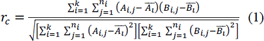

With:

A: scalar vector (first time series)

B: scalar vector (second time series)

n: the number of data in A

k: the number of parts in the current composition for which the correlation is being calculated

n_i_: the number of data in part i

A_i,j_: the jth data (time point) in i^th^ part of vector A

Ai: the arithmetic mean of the i^th^ part of vector A

Since in our data, we did not divide the time series in multiple parts i, the equation was simplified to the following Equation (2):

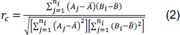

### Protein extraction and analysis

Cell pellets were sampled in the exact same way as for RNA extraction and protein was extracted from the cell pellets as following. Cells were grinded in liquid nitrogen on a pre-frozen mortar and a pestle. Lysis buffer (Sodium Deoxycholate (12 mM) (Wako), Sodium N-dodecanoylsarcosinate (Wako) (12 mM), Ammonium bicarbonate (50 mM) (Nacalai Tesque), Protease Inhibitor Cocktail (25X) (Roche)) was then added to the cells. After 5 rounds of cell grinding, the powder was transferred to a sterile Falcon tube. After thawing the cells on ice, they were transferred to a 1.5 mL sterile Eppendorf tube. The cells were then sonicated using Bioruptor II UCD-200 (20 minutes with intervals of 1 minute). They were then centrifuged at 2,300 x g, 4°C, for 10 minutes. After transferring the supernatants to a new 1.5 mL sterile Eppendorf tube, these were again centrifuged at 13,000 x g, 4°C, for 10 minutes to further remove debris. Protein quantification was assessed using BCA protein assay kit from ThermoScientific following the manufacturer’s instructions. After adjusting the concentration to 1 mg/mL, 1 M of dithiothreitol was added to break down protein disulfide bonds. 1 M of iodoacetamide was added for peptide mapping purposes. After a 5-fold dilution in 50 mM ammonium bicarbonate, the proteins were digested by incubating the samples with 1 µg of Lys-C (Wako) at 37°C for 3 hours, then with 1 µg of trypsin from porcine pancreas (Sigma) at 37°C for 16 hours. After digestion, the sample was acidified with 1% trifluoroacetic acid (TFA) and sonicated for 10 minutes at intervals of 1 minute. The supernatant was then desalted in C18-StageTips, dried under reduced pressure, and stored at −80°C until analysis. The sample was dissolved in 0.1% formic acid 2% ACN and subjected to shotgun proteomic analysis using a nanoElute/timsTOFPro system (Bruker) equipped with a self-packed capillary column (ACQUITY UPLC BEH C18, Waters, 1.7 µm, 75 µm i.d., 250 mm length). The acquired mass spectra were analyzed as described in Proteome analysis part.

### Proteome analysis

Proteomics analysis was carried out as follows. After measuring the mass of the peptides and proteins in our sample, MSFragger (v3.4) handled peptide identification,^56^ and peptide-spectrum validation was done using PeptideProphet and ProteinProphet from Philosopher (v3.4.13 and v4.1.1).^57^ Label-free quantification with match between runs was done using IonQuant (v1.5.5 and v1.7.17) through the FragPipe pipeline v16.0 (for mapping the proteins to database from UniprotKB) and v17.1 (for mapping the proteins to database from the new resulting annotation).^58, 59^ The database used for protein identification was downloaded from UniprotKB (accessed with 2021-09 release) with adding contaminants and false positives.^60^ Downstream analysis was performed using Bioconductor package MS-DAP v1.0.^61^

## Results

### Long-read direct RNA sequencing combined with short-read cDNA sequencing and 5’-CAGE allows reannotation of the transcriptome of fission yeast during glucose starvation

As shown in Figure 1a, fission yeast is exposed to glucose starvation for 120 minutes: after growing in glucose rich medium (6% glucose), the cells are transferred to a medium with only 0.1% glucose during exponential growth, inducing a stress response that is similar to glucose starvation but which does not induce cell death.^62^ Total RNA is extracted at 4 different time points (0 (rich), 15 (early), 60 (mid) and 120 (late) minutes). In order to increase the accuracy of 5’- and 3’-end identification, the transcriptome was sequenced using three methods: (a) long read direct RNA sequencing and (b) shotgun cDNA sequencing (RNA-seq) on poly(A)-enriched RNA, and 5’-CAGE. Using FLAIR and SQANTI3 analysis pipelines to combine these datasets, the fission yeast transcriptome was reannotated taking into account all transcript isoforms during glucose starvation and compared with the reference transcriptome annotation available in Pombase.

**Figure 1.**
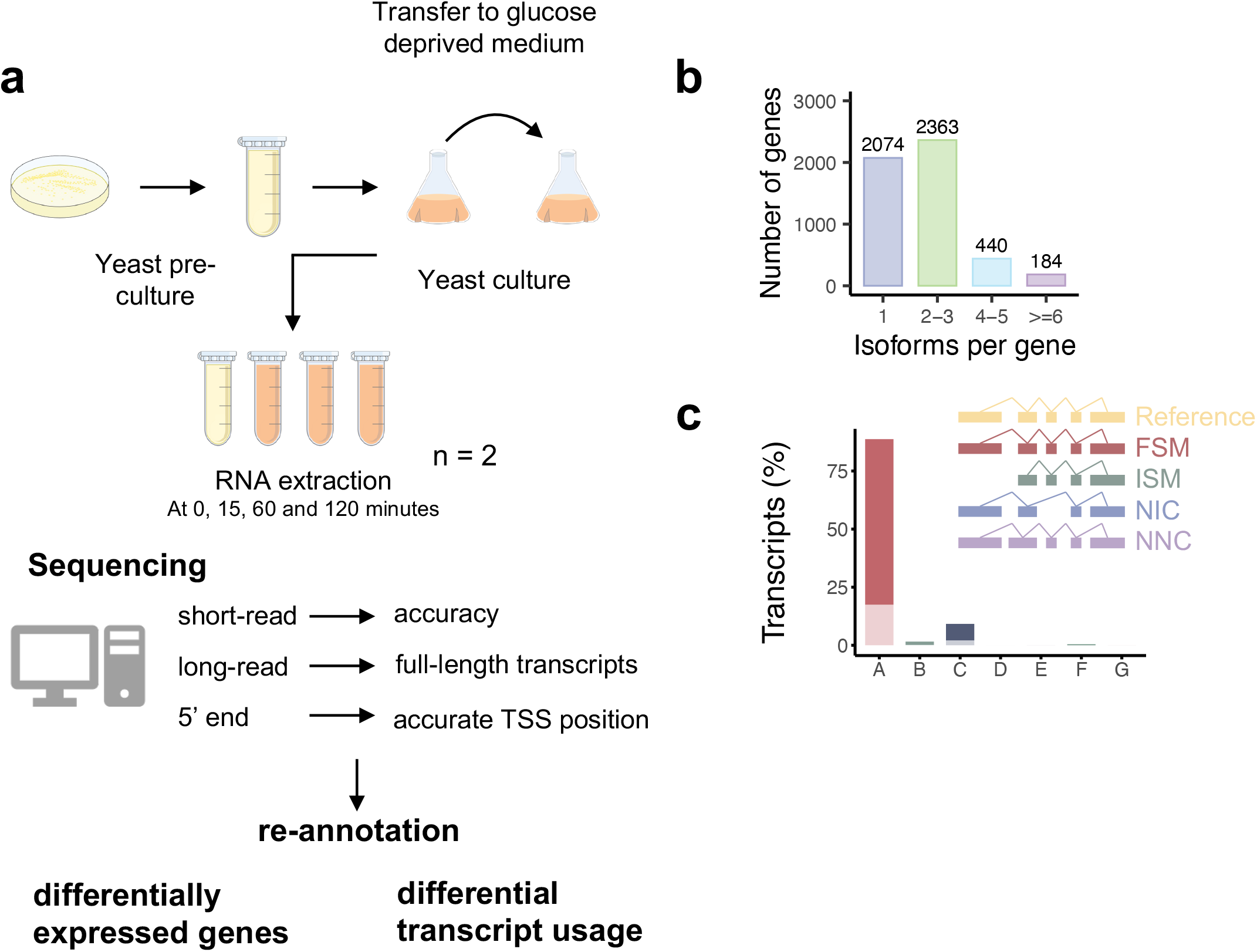
(A) Workflow of the study of fission yeast exposed to glucose starvation with the different sequencing technologies implemented (B) Number of transcript isoforms found per gene (C) Various types of transcripts as classified by SQANTI3 workflow with the percentage of the transcripts belonging to each category A – FSM, B – ISM, C – NIC, D – Genic Genomic, E – Antisense, F – Fusion, G – Intergenic

In this study, the annotation produced by FLAIR/SQANTI3 will be referred as ‘SQANTI’ while the annotation produced by merging the existing Pombase annotation with our newly generated SQANTI annotation will be called ‘PombasePlus’.

The newly identified transcripts are characterized in figures 1b **to** 1d. As shown on Figure 1b, most of the genes had between 1 or 2-3 isoforms (41.0% and 46.7% respectively) and a small portion of genes had more than 4 isoforms (**Supplementary Table 1**) (12.3% (624/5061)). When compared to the reference, SQANTI3 categorizes the transcripts into various categories: Full Splice Match (FSM), Incomplete Splice Match (ISM), Novel In Catalog (NIC) and Not Novel in Catalog (NNC). ISM are isoforms that ‘have fewer 5’ exons than the reference, but each internal junction agrees. The exact 5’ start and 3’ end can differ by any amount’. NIC transcripts are defined as an isoform that ‘does not have a FSM or ISM, but is using a combination of known donor/acceptor sites’ (citation from the SQANTI user manual).^40, 42^ As observed on Figure 1c, most of the transcripts in the annotation produced by the pipelines are FSM (9,839 (88.7%) in brown), 1,020 (9.2%) are NIC, and 175 (1.6%) are ISM.

We also defined the overlap between coding and non-coding transcripts found in both SQANTI and Pombase’s annotation. Moreover, 38 genes detected as fusion genes were considered artefacts and removed before further analysis. The coding potential of genes was assessed by GeneMark from the SQANTI3 workflow. Coding genes, which represent 77.5% of all the genes, found in both datasets. A small portion of the coding genes are newly found by SQANTI (121 (2.30%)). After mapping these genes, we found that they map to both a coding and a non-coding gene which is annotated as ‘predicted untranslated region (UTR)’. However, the overlap was smaller for non-coding genes: only 627 (7.67%) are found in both SQANTI and Pombase. Some of the genes not found in the overlap are genes that were erroneously classified as non-coding by GeneMark based on the small size of the ORF, despite being categorized as coding in Pombase and experimentally validated. Among these genes, there are also unlisted antisense genes, which were (**Supplementary Table 1**) opposite of *alp31+* (SPAC8E11.07c), *mam2+* (SPAC11H11.04) and *erg4+* (SPAC20G4.07c). Genes that were only present in Pombase’s annotation include genes that were described in previous literature and may also include genes giving birth to non-capped and/or non-polyadenylated transcripts. Hence, Figure 2a shows that our reannotation obtained with the FLAIR/SQANTI3 pipelines is for the most part consistent with the existing Pombase annotation in terms of gene content, with a number of novel coding and non-coding transcripts that correspond to 7.6% and 2.4% of their known annotated transcripts respectively. The list of all transcripts and their respective genes are found in **Supplementary Table 2.** The gene content is consistent with the previous annotation, but our pipeline’s purpose is to provide new insights on the precise 5’- and 3’-ends of transcripts, the results of which are described in the following sections.

**Figure 2.**
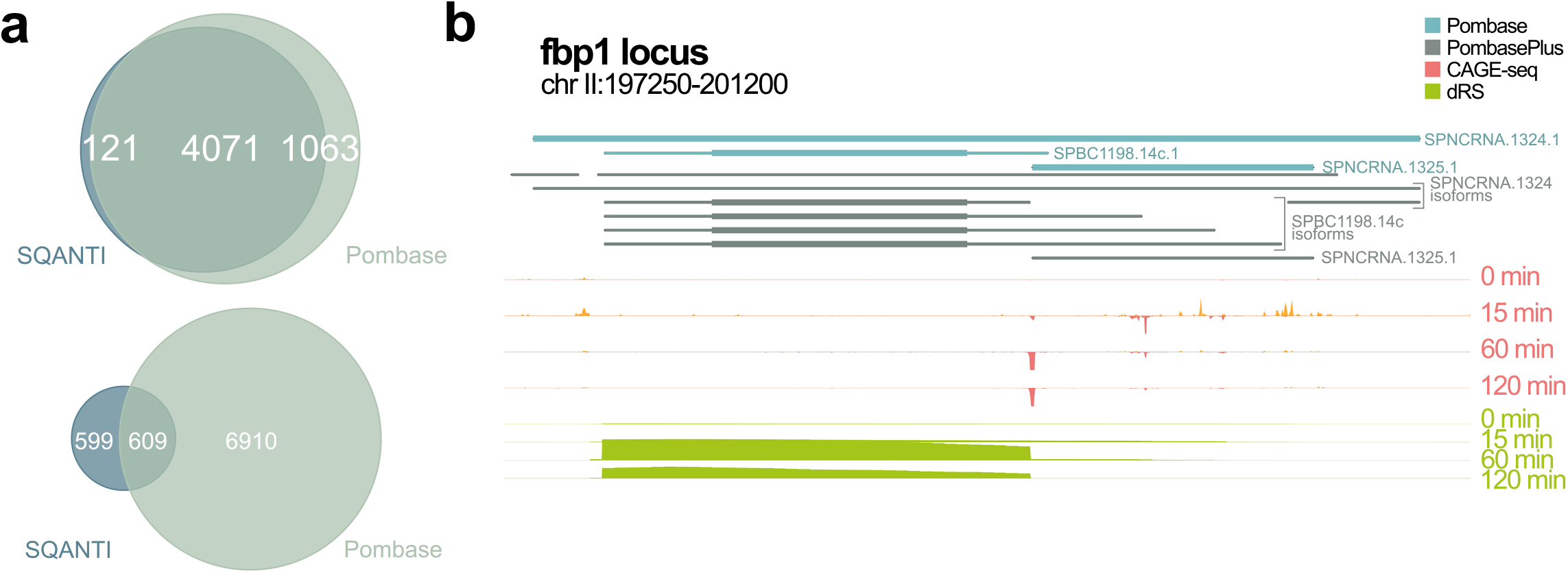
(A) Overlap of coding genes between SQANTI and Pombase and overlap of non-coding genes between SQANTI and Pombase (C) Pombase and PombasePlus annotation and Next Generation Sequencing (NGS) tracks at *fbp1*+ locus

### Differential gene expression shows an upregulation in carbon metabolism associated with a downregulation in ribosome biogenesis

In response to glucose starvation, the fission yeast shifts its metabolism by inducing or repressing the gene expression.

Figure 3a shows the evolution of gene expression patterns during glucose starvation that were observed among 561 significantly differentially expressed genes (DEGs) which were defined as genes with a significantly different expression level at least one time point compared to the glucose-rich condition (0 minutes) (|log2(Fold Change)| ≥ 2, p-value ≤ 0.05, Wald test). These DEGs represent 5.7% of the gene set in our new PombasePlus annotation (9,684 genes). More DEGs are observed at 60 minutes (424 (upregulated:downregulated) (251:173) genes compared to 265 (188:77) and 172 (123:49) genes for 15 and 120 minutes respectively), suggesting that although some genes are induced or repressed earlier on, the stress response induced the biggest change in terms of number of DEGs at 60 minutes into glucose starvation.

**Figure 3.**
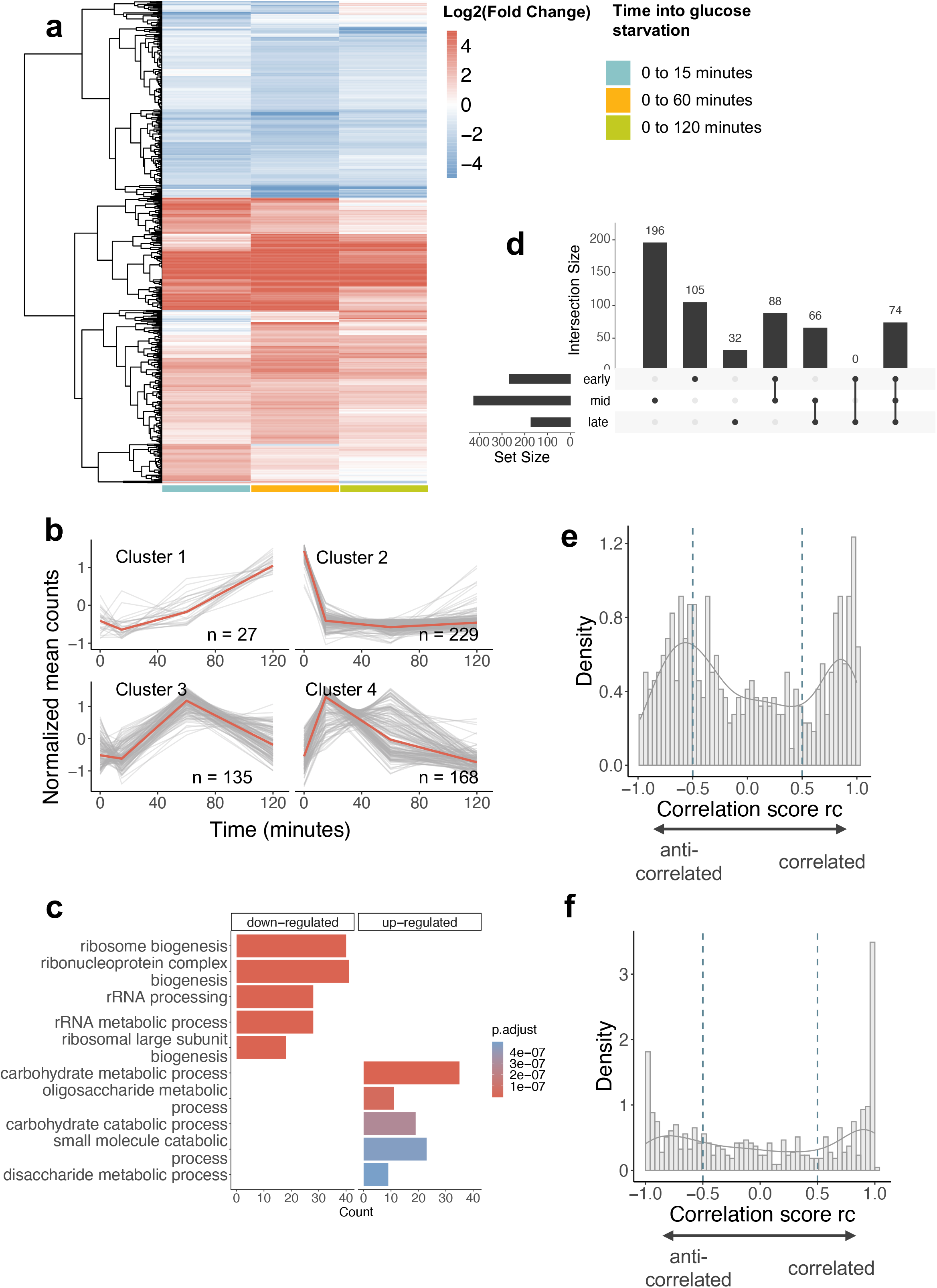
(A) Heatmap of gene expression of the differentially expressed genes (DEGs) during glucose starvation compared to glucose rich condition (cutoffs: |log2(FC)| > 2, *p*-adj < 0.05, n = 2) (B) KEGG enrichment pathway for downregulated and upregulated DEGs ranked by *p*-adj (C) Upset plot showing the overlaps between the DEGs found at each stage into glucose starvation (early: 15 minutes, mid: 60 minutes, late: 120 minutes). **Supplementary Table 4** shows the different genes in each category, going from left to right. (D) Clustering of gene expression of the DEGs during glucose starvation (E) Correlation score rc distribution for pairs of DEG with their antisense (with no lag) (F) Correlation score rc distribution for pairs of DEG with their antisense (with lag).

Hierarchical clustering on the DEG’s expression during glucose starvation revealed 4 clusters: cluster 1 (permanent induction) with 27 genes, cluster 2 (permanent repression) with 229, cluster 3 (mid-transient repression) with 135 and cluster 4 (early-transient induction) with 168 genes (Figure 3b and **Supplementary Table 3**). KEGG pathway enrichment analysis (Figure 3c) on both upregulated and downregulated genes at all time points (False Discovery Rate (FDR) < 0.05) showed that upregulated genes tend to be involved in carbohydrate metabolic process and oxidoreductase activity. On the other hand, downregulated genes are involved in ribosome biogenesis, rRNA processing and ribonucleoprotein complex biogenesis, consistent with previously observed transient global translation inhibition observed in glucose starvation.^13, 63^ It is likely that the fission yeast reallocates some of the resources that were spent on ribosome biogenesis for use in carbon metabolism to ensure survival during the shift. The KEGG pathway enrichment changes dynamically over the course of the response, consistent with previously described resuming of translation and cell division at 120 minutes (**Supplementary Table in Supplementary 2**).

When comparing the overlap of DEGs between the different stages into glucose starvation (early, mid and late) (Figure 3d and **Supplementary Table 4**), early and mid-stages overlap by 88 genes while mid and late overlap by 66 genes. Over the course of the glucose starvation response, 74 genes are constantly differentially expressed compared to glucose-rich. In contrast, a higher number of genes were differentially expressed at only one stage (105, 196 and 32 genes respectively for 15, 60 and 120 minutes), in other words showing a transient expression change. Supplementary Figure 1 also shows that the global correlation coefficient between RNA expression at 15 minutes and 60 minutes (r^2^ = 0.85) is lower than the one between 60 minutes and 120 minutes (r^2^ = 0.94), suggesting that although the number of DEGs itself was the highest at 60 minutes, the overall amplitude of the change was biggest at 15 minutes.

### A subset of differential expressed genes is associated with antisense transcription on the opposite strand

Previous studies on the *fbp1+* gene showed that its expression in glucose starvation is anticorrelated with the expression of antisense RNA on the opposite strand. Our new annotation leveraging full-length RNA sequencing allowed us to identify all four previously sense-strand overlapping transcripts as well as three antisense RNA isoforms on the opposite strand, one of which is novel and overlapping with the 3’-end of sense transcript but not with the ORF.^14^ We found 538 pairs of DEG-antisense, with some DEGs associated with multiple antisense transcripts based on the annotation file. Among these, 469 unique DEGs out of the total 561 were associated with at least one antisense (83.6 %). This was not so different from the genome-wide value: 8,378 expressed genes out of 9,716 which had an antisense (86.2 %), suggesting that DEGs are not particularly enriched in antisense annotations. To better understand the mechanisms, we investigated the potential co-expression between these pairs.

To determine whether a pair of sense-antisense is correlated or anticorrelated, a compositional correlation score rc was calculated (for details, cf Material and Methods section), with rc < −0.5 indicates anticorrelation and rc > 0.5 correlation. Figure 3e shows the distribution of the rc for these pairs. The number of DEGs that are either positively or negatively correlated with their antisense are roughly equal in size (168 and 160 respectively) when no time lag is considered.

Next, we hypothesized that the antisense gene transcription may temporally precede the sense transcription, as suggested by previous studies showing a regulatory effect of the antisense on the remodeling of the chromatin structure in *cis*.^64, 65^ However, past studies in fission yeast do not take into a possible delayed effect when studying correlation. To investigate this, the rc with the DEG’s expression at time points 15, 60 and 120 minutes was calculated as an input for a scalar vector A, and at time points 0, 15 and 60 minutes for the associated antisense gene as an input for a second scalar vector B. The distribution of the rc for the same pairs at lagged time points in the glucose starvation is shown on Figure 3f.

Table 1 shows that when considering a time lag, significantly more strongly anticorrelated gene pairs (60 genes with rc ≤ −0.9) were found than when comparing simultaneous time points (19 genes with rc ≤ −0.9 for no lag). This suggests that for anticorrelated sense-antisense pairs, expression of the antisense does indeed tend to precede the sense’s expression. Interestingly, a population of positively correlated sense-antisense pairs were detected in roughly equivalent numbers, suggesting that some opposite-strand pairs of genes may operate using a different mechanism. Correlation scores rc for each pair of DEG-antisense, both non-lag and lag, are presented in **Supplementary Table 5** along with their expression levels.

**Table 1.** Number of gene pairs with an absolute correlation score more than 0.5 and 0.9

|  |  |  |
| --- | --- | --- |
| **no lag** | $\leq -0.5$ | 160 |
| | $\geq 0.5$ | 168 |
| **lag** | $\leq -0.5$ | 179 |
| | $\geq 0.5$ | 141 |
| **no lag** | $\leq -0.9$ | 19 |
| | $\geq 0.9$ | 65 |
| **lag** | $\leq -0.9$ | 60 |
| | $\geq 0.9$ | 69 |

When testing for antisense transcription preceding that of sense transcription, 25.1% of all DEG-antisense pairs are positively correlated, and 31.9% are anticorrelated. The former are likely to operate by a different mechanism than the latter.

### Differential transcript usage induces protein domain gain and loss during glucose starvation

In addition to differential gene expression, it has been previously reported that fission yeast uses alternative transcription start sites (aTSS) in other stress responses (nitrogen depletion, heat shock or oxidative stress).^10^ Our analysis reveals 322 differential transcript usage events (DTUs), defined by alternative 5’- and 3’-ends but not necessarily any change in gene expression levels.

When comparing the overlap between all DEGs (561) with the DTUs identified here, an overlap of 36 genes was observed (Figure 4a). The relatively low overlap between these two sets of genes suggests that that fission yeast regulates its metabolism in a bimodal manner (**Supplementary Table 6** shows the DEG and DTUs with the overlap). Genes with isoform switches have various consequences, which are investigated below.

**Figure 4.**
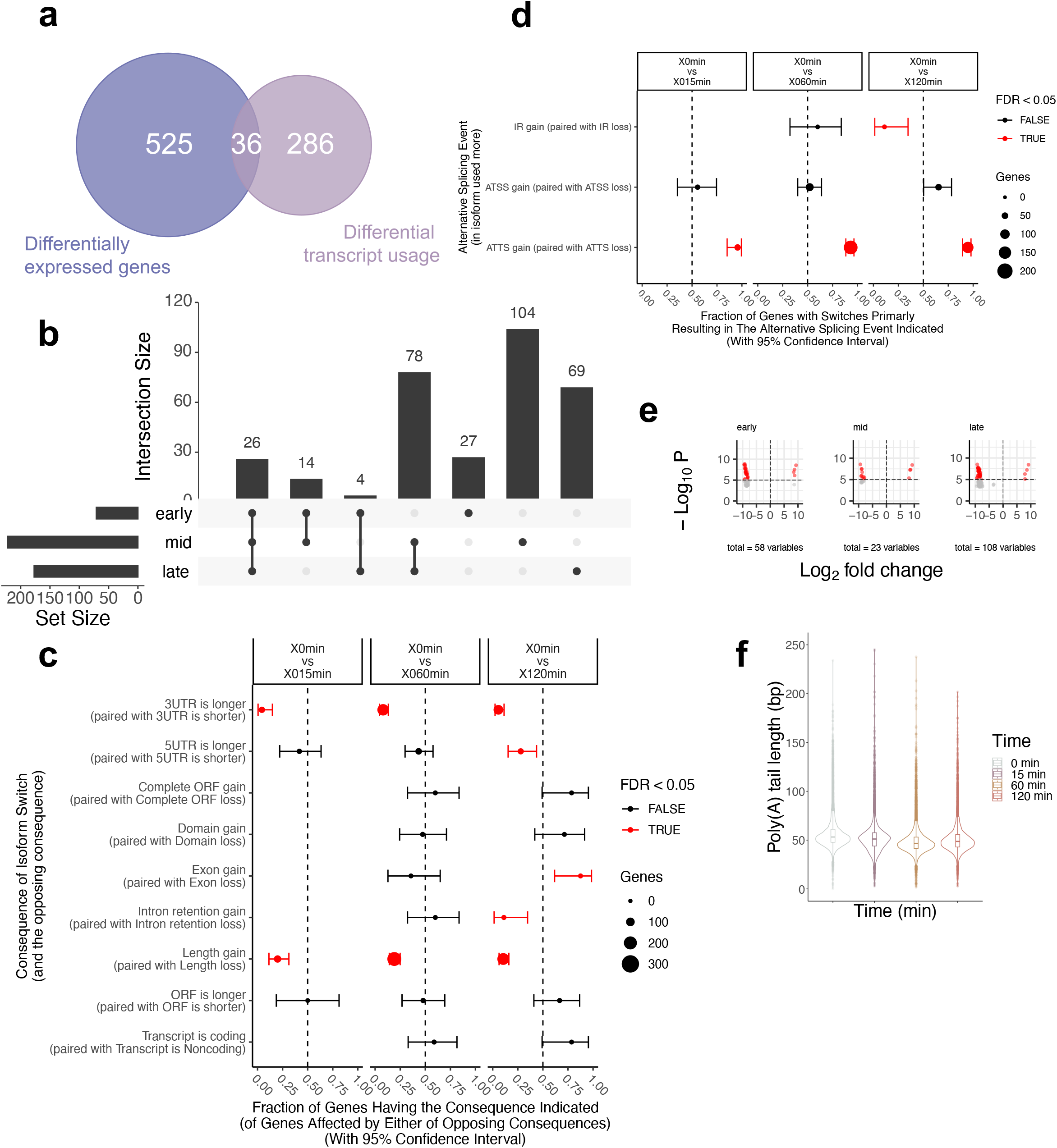
(A) Venn diagram showing the overlap between DEG and DTU (B) Upset plot showing the overlaps between the DTUs found at each stage into glucose starvation (early: 15 minutes, mid: 60 minutes, late: 120 minutes). **Supplementary Table 7** shows the different genes in each category, going from left to right. (C) Fraction of genes having the consequence indicated when compared to glucose rich condition (D) Fraction of genes having the consequence indicated when compared to glucose rich condition (E) Violin plots of the distribution the poly(A) tail lengths at each time into glucose starvation for each gene

Like for the DEGs, the overlap between the DTUs found in the early and late stages is minimal (4 genes) compared to the overlap between the other time points: the early-mid and mid-late overlapped by 14 and 78 genes respectively. 26 genes are consistently identified as a DTU compared to the glucose rich condition (**Supplementary Table 7**). Similarly, distinct sets of genes are DTUs at each time point during glucose starvation, again consistent with dynamic regulation within the first two hours of glucose starvation.

Associated with the various alternative isoform switch events, changes in functional consequences happen. Transcripts are significantly shorter with a length loss in 3’-UTRs at all time points during the stress (for 44, 151 and 117 genes at early, mid and late stages respectively) and in 5’-UTRs (for 32 genes) later into glucose starvation (120 minutes). One advantage of DRS is the possibility to assess the length of the poly(A) tail of each transcript. A significant shortening of the poly(A) tail length is observed as well, though with some genes had a longer poly(A) tail (SPCC1919.12c, SPNCRNA.1223, SPNCRNA.4537, SPNCRNA.964 at all time points, and SPNCRNA.3213 only at 15 minutes) (Figures 4e and 4f). We used the Wilcoxon signed-rank test to compare the poly(A) tail length at each time point with 0 minutes as decribed in the section Material and Methods. The *p-*values are extremely small for all time points compared to glucose-rich, indicating that the poly(A) tail length is significantly smaller at each time point in glucose starvation (Table 2) [Table 2 **near here**]. Overall, there is a global shortening of the poly(A) tail length. We also noticed that genes with a shorter 3’ UTR and genes with a shorter poly(A) tail length did not overlap.

**Table 2.** Related t-test results comparing poly(A) tail lengths at different time points during glucose starvation

|  | **Statistic value** | ***p*-value** |
| --- | --- | --- |
| **15 min vs 0min** | 15257928.0 | 1.65E-224 |
| **60 min vs 0 min** | 9782070.0 | 0.0 |
| **120 min vs 0 min** | 11384537.0 | 0.0 |

Among the 9,888 genes that have a poly(A) tail at any time point, 122 genes have a poly(A) tail with a significant change in length. Interestingly, 89.3% of these genes were non-coding (142/159) with one among the coding genes being dubious (**Supplementary Table 8**). The changes in poly(A) tail length when comparing 60 minutes to glucose rich are less prominent compared to the changes at 15 and 120 minutes.

Associated with shorter 5’ and 3’ UTRs, domains gains and losses are also discovered for a small set of genes (15 and 12 domains respectively). The gained and lost domains detected at each time point are described in **Supplementary Table 9**. These domains were predicted computationally by Pfam and we quantified the peptides from our proteomics data (cf Material and Methods) at the domain level to validate or the gain and/or loss as presented in **Supplementary Table 10.** Domains that were gained during glucose starvation by isoform switching include domains involved in the transcriptional activity of RNA polymerase II and transcription elongation regulation, vegetative growth, spore formation and mitophagy. Domain losses were observed on proteins involved in sugar transport and ribosome subunits such as ribosomal proteins L29, L35ae, and S17, which is consistent with the downregulation in ribosome biogenesis described in previous section.

### Upregulated differential detected proteins are also involved in carbon metabolism and downregulated ones are involved in endocytosis, promoting the fission yeast cells’ survival

The effect of glucose starvation on the proteome landscape of the fission yeast was studied at the same 4 time points. In contrast with the results of transcriptome analysis, where induced transcripts tend to be expressed at a basal level in glucose-rich, many proteins were not detected at all in glucose rich but specifically produced later in the stress response. The z-score calculated by MS-DAP for differential detection was the basis for our analysis. Figure 5a shows differential z-score patterns during glucose starvation for 379 proteins with a |z-score| > 2 at any time point into the stress, defined as differentially detected proteins (DDPs). This represents approximately 7.4% of all the proteins listed in UniProt database for the fission yeast (out of 4,021 proteins detected). Interestingly, compared to RNA expression, we see that there are more DDPs at early stages compared to mid stages (217 (upregulated:downregulated) (95:122) proteins at early step, 198 (145:53) at mid and 217 (177:40) at later stages). While there are more DDPs at 15 minutes, 56.2% (122/217) are downregulated, later stages into the stress have a reversed ratio of upregulated to downregulated proteins, with upregulated ones being more dominant.

**Figure 5.**
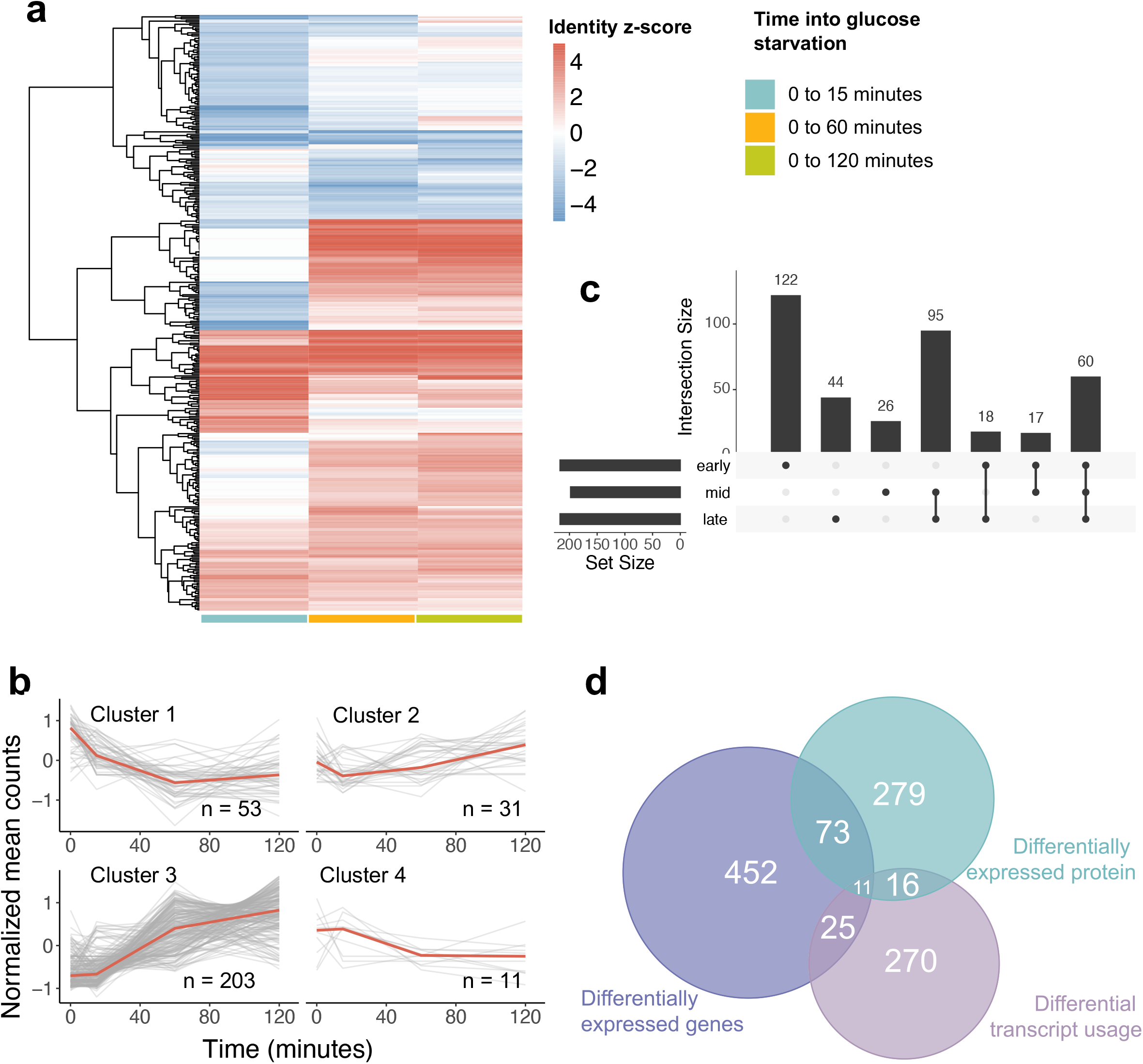
(A) Heatmap of protein detection using z-score of differentially detected proteins (DDPs) during glucose starvation compared to glucose rich condition (cutoffs: |z-score| > 2, n = 4) (B) Clustering of the protein intensity of differentially expressed proteins during glucose starvation (C) Upset plot showing the overlaps between the DDPs found at each stage into glucose starvation (early: 15 minutes, mid: 60 minutes, late: 120 minutes) **Supplementary Table 12** shows the different genes in each category, going from left to right. (D) Venn diagram showing the overlaps between DEGs, DTUs and DDPs.

KEGG pathway enrichment analysis on upregulated proteins shows an enrichment in proteins involved in the TCA cycle, 2-oxocarboxylic acid metabolism, biosynthesis of amino acids and carbon metabolism, while the downregulated proteins are involved in endocytosis, purine metabolism, nucleocytoplasmic transport and spliceosome. Similarly, we clustered the DDPs’ intensity during glucose starvation into 4 clusters, cluster 1 (permanent repression) with 53 proteins, cluster 2 (early transient repression) with 31, cluster 3 (permanent induction) with 203 and cluster 4 (early transient repression) with 11 proteins (Figure 5b). The list of proteins in each cluster is presented in **Supplementary Table 11**. Cluster 2 is significantly enriched in Protein K11-linked ubiquitination, positive regulation of autophagy, Anaphase-promoting complex-dependent catabolic process and Golgi to plasma membrane transport for biological processes.

We also compared the overlap of DDPs at different stages into glucose starvation (Figure 5c and **Supplementary Table 12**). As observed previously, the mid and late stages have the biggest overlap (95 proteins), while early/mid and early/late have a smaller one (17 and 18 proteins, respectively). This suggests that the proteome changes most dynamically in the early stage (15 minutes). Likewise, Figure 5d shows the overlap between DEGs, DTUs and DDPs at all time points into glucose starvation. The overlap between DEGs and DDPs is 15.0% of all DDPs and that between DTUs and DDPs is 8.4% of all DTUs, confirming previous results showing that the correlation between gene expression and protein expression is low (Supplementary Figure 3). Hence, both DEGs and DTUs are partially involved in the changes in the translational response. DEGs only are enriched in proline transmembrane transport, proline import across plasma membrane, trehalose metabolic process, cellular response to heat and disaccharide metabolic process. For the DTU, these are enriched in membrane protein complex, golgi apparatus, organelle component, organelle membrane and endomembrane system when doing gene ontology analysis on cellular component. Figure 6 summarizes the major metabolic pathways with different genes involved and their gene and protein expression during glucose starvation. Most DEGs and DDPs are involved in starch and sucrose, galactose, glycolysis, fructose, pentose phosphate, TCA cycle and tyrosine metabolism, with both DEGs and DDPs involved in starch and sucrose and galactose metabolisms upregulated during glucose starvation, with a lag for DDPs where they are downregulated first.

**Figure 6.**
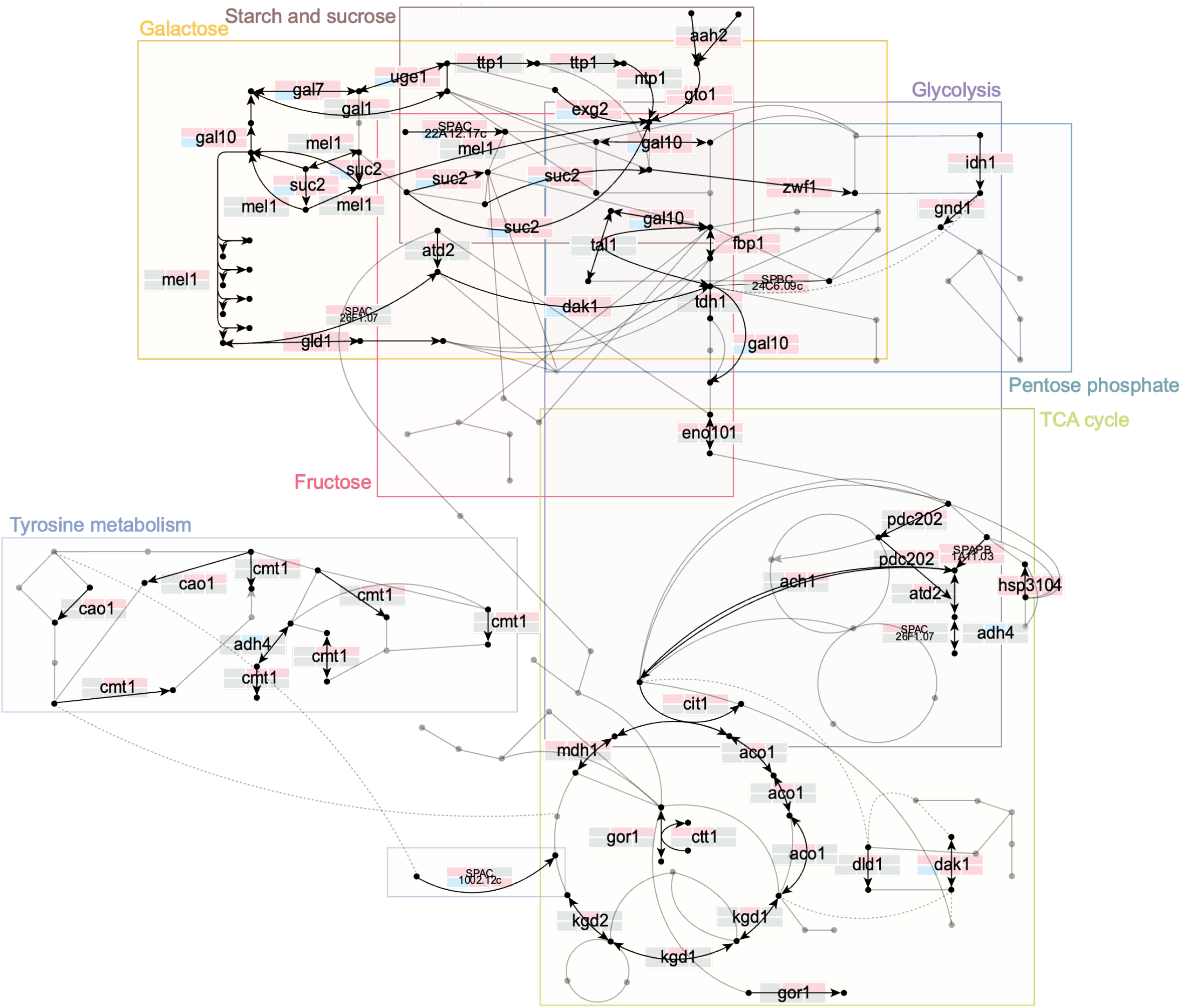
DEGs and their gene and protein expression during glucose starvation and the metabolic pathways they are involved in. Each arrow corresponds to a reaction. Each gene producing the protein associated is written on the figure if expressed at any time point or any transcript or protein level. For each gene, their expression during glucose starvation is shown at each time point (15, 60 and 120 minutes compared to 0 minutes). The first row is the expression at the gene level while the second row is the expression at the protein level. If the expression is upregulated compared to 0 minutes, the rectangle is red. If it is downregulated, it is blue. If it is grey, it is not expressed at this specific time point.

## Discussion

In the past decades, new sequencing technologies have emerged and improved allowing the discovery of new mechanisms of gene regulation. In this study we focus on identifying differential 5’- and 3’-transcript in glucose starvation in fission yeast.

As a result, differentially expressed genes during glucose starvation were classified into 4 clusters depending on the timing of their activation. The Gene Ontology results for these correspond well to previous results described in Oda *et al*.^14^, therefore confirming that our reannotation, while providing much more detail at the transcript isoform level, is largely consistent with past literature at the gene level.

Our study, by allowing us to precisely define the boundaries of overlapping transcripts, also allowed us to identify DEG-antisense pairs at the isoform level in greater detail. These include known DEG-antisense isoforms in glucose starvation that have been previously confirmed by Northern blotting.^14, 15^ As Muskovic *et al*.^66^ previously suggested, the transcription of regulatory lncRNAs precedes the changes in the sense gene expression, we showed that in the fission yeast, the correlation between the expression of DEGs and their antisense’s is more highly correlated with a time delay into glucose starvation, suggesting that the antisense genes are transcribed first. We also found that antisense transcription concerns around 86.2% of genes. Wery *et al*.^67^ found a similar percentage of 76.9% in fission yeast. A slight discrepancy of these percentages can be explained by updates in the Pombase database since 2018 (date of publication of Wery *et al*.’s study) and the fact that they are using different sequencing methods, in different growth conditions. In addition, we found two populations of pairs of antisense-sense genes, one with anticorrelated gene expression (31.9%) and the other one with correlated gene expression (25.1%). Moreover, previous studies that describe anticorrelation as their main finding base their conclusions on a global trend using Spearman rank correlation between antisense and sense expression variables and did not expose the fission yeast cells to any stress,^67, 68^ explaining the difference with our results. Previous studies also show the existence of both positively and negatively correlated antisense-sense pairs,^69^ with divergent pairs of antisense-sense being anticorrelated and the convergent ones being positively correlated. While our study shows two different populations of antisense-sense pairs as Leong *et al* found, we did not find a clear relationship between the divergence of these pairs and their correlation score. In conclusion, we observed both a positive and negative correlation in antisense-sense pairs’ gene expression during glucose starvation in a time lag. This aspect is relatively novel as previous studies have not considered the time lag when assessing correlation.

Following this, the precise quantification of overlapping transcripts unveils aTSS and aTTS switches happening during glucose starvation which have not been described before in the fission yeast. Globally, we observed significant shortening of the 3’-UTRs, alongside with poly(A) tail length from early stages into glucose starvation. 5’-UTRs are significantly shorter at later stages. Studies in other organisms exposed to several stresses have shown changes in 3’-UTRs and poly(A) tail lengths mainly.^70–75^ 3’ UTR shortening and lengthening studies are still incomplete, however, studies in humans and mouse cells suggest that a shortening increases mRNA stability by limiting the access to mRNA degradation mechanisms (such as ribosome binding proteins or miRNA).^70,76–78^

As opposed to genes with a shorter 3’-UTRs, genes with a shorter poly(A) tail length are mainly non-coding genes. Unlike mRNAs, the presence of a poly(A) tail on non-coding RNAs is a signal for the degradation of the transcript.^79^ While past research has investigated the impact of poly(A) tail length on the mRNA, including its stability,^80^ its expression,^81^ and its translational efficiency,^82^ no previous research has been done to describe the impact of the changes in poly(A) tail length on non-coding RNA. Since mRNA with shorter poly(A) tails are targeted for degradation, we hypothesize that the shortening of the poly(A) tails may also be signal for degradation of the non-coding RNAs during glucose starvation. Further investigations are necessary to decipher the mechanism by which these specific non-coding transcripts are targeted. These switches directly impact the length of both the ORFs and the UTRs, inducing changes in proteins domains for 32 genes and 52 different protein domains (**Supplementary Table 9**) involved in transcriptional activity and mitophagy and were validated by proteome analysis. Zheng *et al*. showed in a previous study that complete starvation of the yeast induces mitochondrial fragmentation.^83^ While previous studies have inquired into protein domain changes (gain and loss) between various organisms,^84–86^ our research is the first one to describe protein domain changes during glucose starvation induced by aTSS and aTTS.

On a translation level, as observed in previous studies in various organisms and confirmed by our results (Supplementary Figure 3),^87–89^ correlation between gene expression and protein detection is poor. The use of different technologies and methodologies for detecting the transcripts counts and the protein abundance,^90^ on a technical level, and a time delay between mRNA and protein synthesis, with their stability and half-life being distinct, on a biological level, explain this poor correlation. Both DEGs and DTUs are partially overlapping with DDPs, implying that two different mechanisms are involved in the post-transcription response regulation. DEGs, with their changing expression directly impact changing protein abundance. DTUs, while having their boundaries changed, their expression is not, which indicates that changes in protein abundance might involve other mechanisms such as RNA half-life, protein half-life or the ribosomal activity for these transcripts. Globally, in early stages into glucose starvation, there is a downregulation of proteins, notably permeases and transporter proteins, with an upregulation in proteins involved in ubiquitination and autophagy, suggesting a rapid protease activity in early stages of the glucose starvation, by the recognition of ubiquitinated substrates, which are then digested into peptides.^91^ For confirming these results, additional analysis involving measurements of ubiquitination levels and enzymatic activity of the proteases could be done.

In conclusion, by comprehensively reannotating the 5’- and 3’-boundaries transcript isoforms that are both 5’-capped and polyadenylated in glucose starvation using a combination of three different sequencing methods, our study unveils that both changes in gene expression (DEGs) and shifting of transcript boundaries (DTUs) contribute to the fission yeast response to glucose starvation, with functional consequences at the protein level in terms of domain gain/loss. Our data also shows that the anticorrelation observed at sense-antisense pairs tends to be stronger when a time lag is taken into consideration, and that the poly(A) tails of many non-coding RNAs are shortened in glucose starvation. On messenger RNA, 3’-UTRs shortening following by 5’-UTR shortening highlights the need for in-depth follow-up studies to evaluate the impact of glucose starvation on ribosomal activity, RNA stability, and translation efficiency.

## Fundings

This work was supported, in part, by research funds from the Yamagata Prefectural Government and from Tsuruoka City, Japan. In addition, financial support was provided to LCN by the Shonan-Fujisawa Campus (SFC) Academic Society in 2020 through the Emergency Fund for COVID-19 and the Mori Memorial Fund, as well as the Japan Science and Technology Agency (JST) SPRING project under grant numbers F21660A and F22660A. The early stages of the project were partially funded by the Nestlé Nutrition Council of Japan to JG under grant number NNCJ2016.

## Conflicts of Interest

The authors report there are no competing interests to declare.

## Data Availibity

The data that supports the findings of this study is deposited in Sequence Read Archive (SRA) database under the BioProject <u>PRJNA990323</u> (shotgun cDNA-sequencing, long-read sequencing and CAGE-sequencing data) and in MassIVE under ProteomeXchange dataset <u>PXD043446</u>.

## Supporting information

Supplementary Tables description

Annotation file produced in this study

---

**Supplementary Figure 1.**
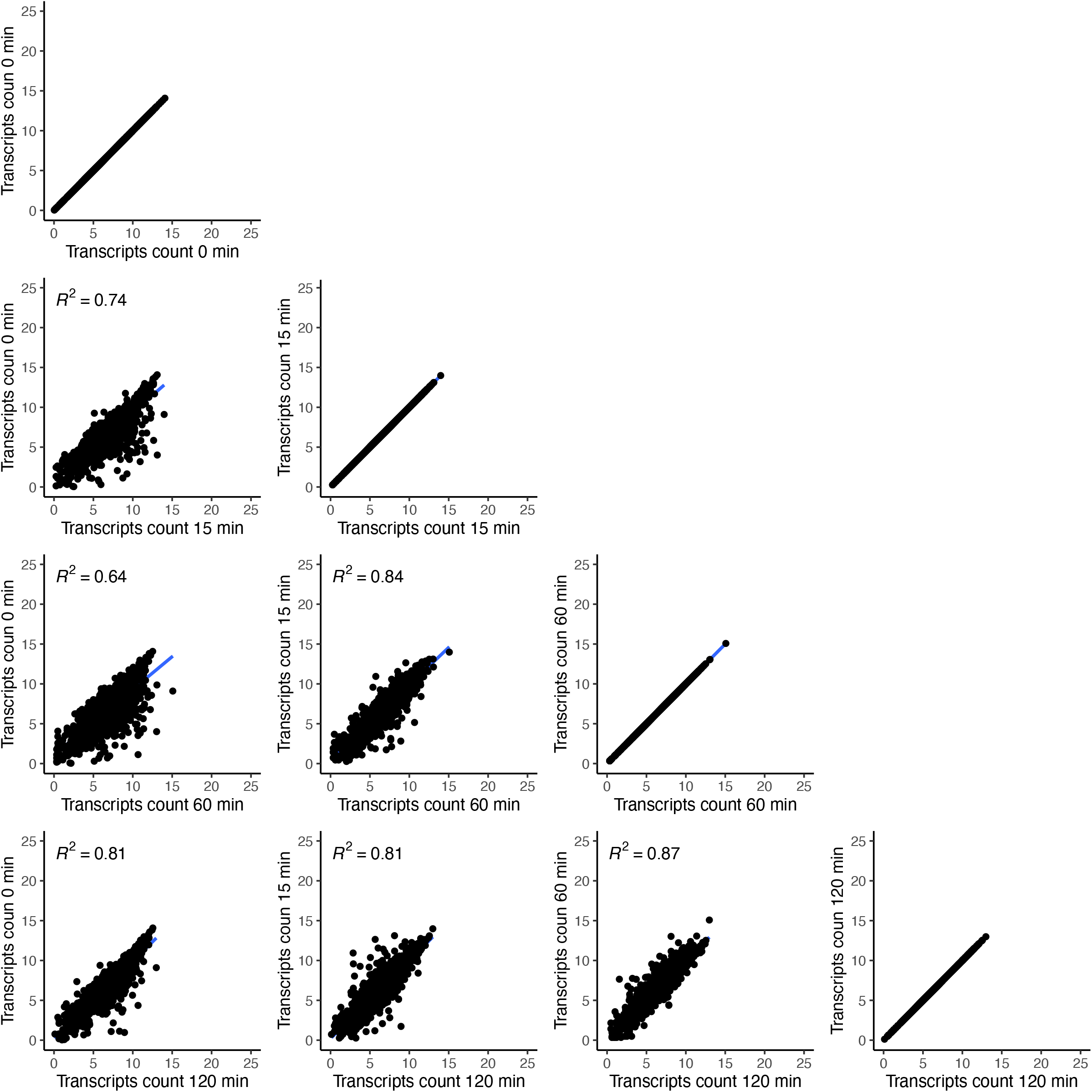
Correlation between RNA expression at each time point

**Supplementary Figure 2.**
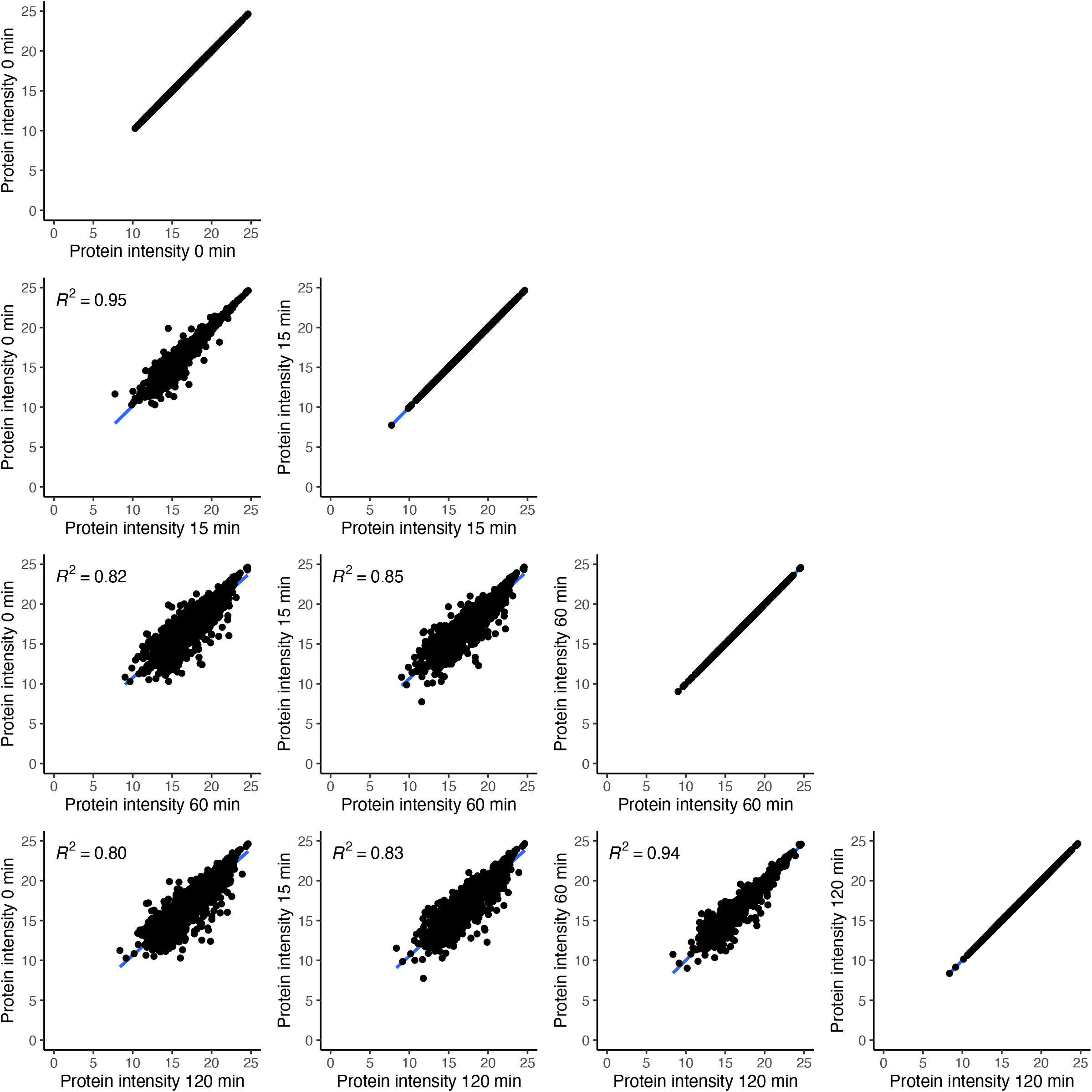
Correlation between protein intensity at each time point

**Supplementary Figure 3.**
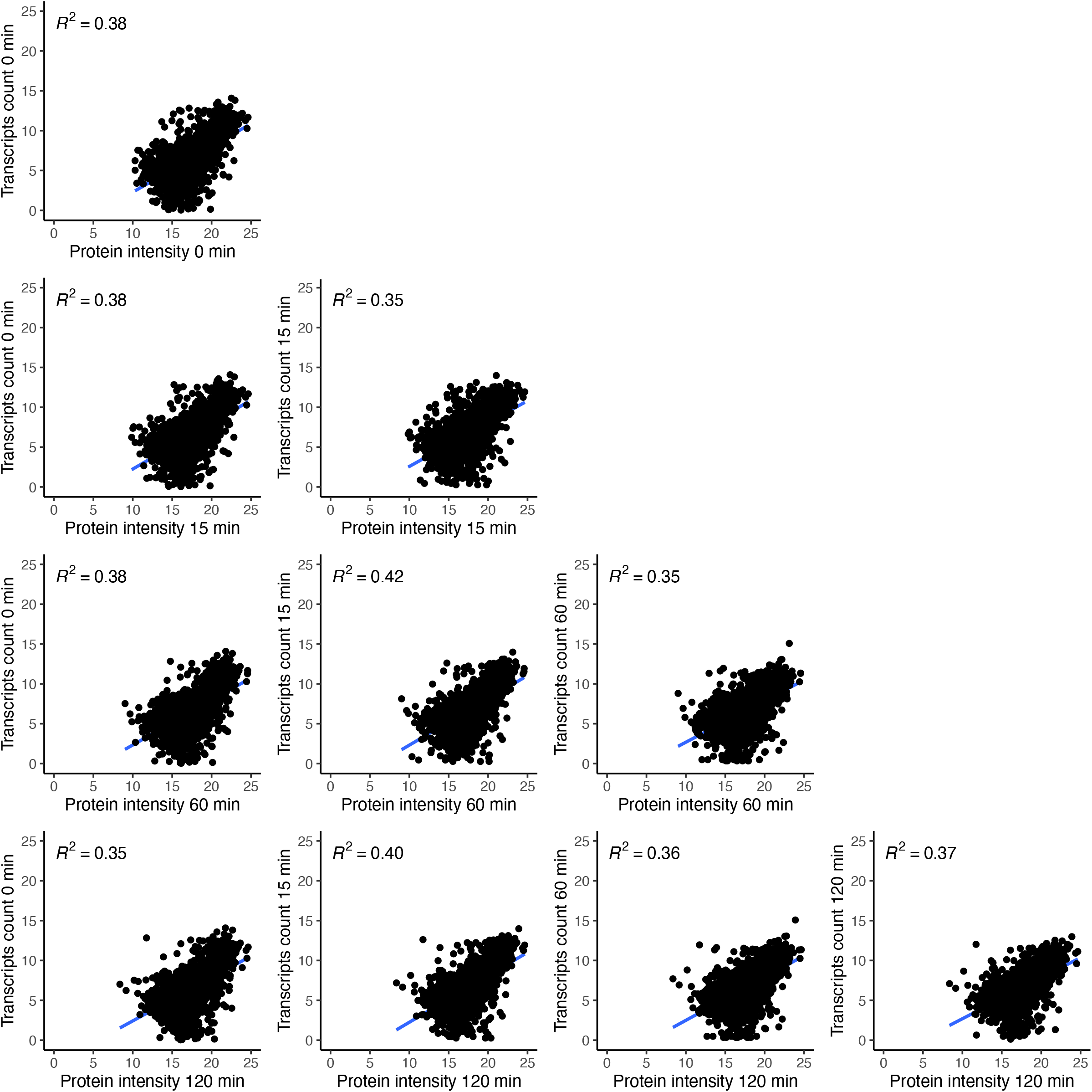
Correlation between RNA expression and protein intensity at each time point

## Notes

https://www.ncbi.nlm.nih.gov/sra/PRJNA990323

https://massive.ucsd.edu/ProteoSAFe/dataset.jsp?task=2c7c71ab07644cb29c61ca561cb845cc

### Competing Interest Statement

The authors have declared no competing interest.

### Summary of Updates

The abstract was re-written both for clarity and to fit within the 200-word requirement for our target journal. The funding section was updated to include more accurate information.

## References

1. Mlekusch W, Lamprecht M, Öttl K, Tillian M, Reibnegger G. A glucose-rich diet shortens longevity of mice. Mechanisms of Ageing and Development. 1996;92(1):43–51. doi:10.1016/S0047-6374(96)01801-5

2. Liu Z, Jia X, Duan Y, Xiao H, Sundqvist K-G, Permert J, Wang F. Excess glucose induces hypoxia-inducible factor-1α in pancreatic cancer cells and stimulates glucose metabolism and cell migration. Cancer Biology & Therapy. 2013;14(5):428–435. doi:10.4161/cbt.23786

3. Masuda F, Ishii M, Mori A, Uehara L, Yanagida M, Takeda K, Saitoh S. Glucose restriction induces transient G2 cell cycle arrest extending cellular chronological lifespan. Scientific Reports. 2016;6(1):19629. doi:10.1038/srep19629

4. Zou K, Rouskin S, Dervishi K, McCormick MA, Sasikumar A, Deng C, Chen Z, Kaeberlein M, Brem RB, Polymenis M, et al. Life span extension by glucose restriction is abrogated by methionine supplementation: Cross-talk between glucose and methionine and implication of methionine as a key regulator of life span. Science Advances. 2020;6(32):eaba1306. doi:10.1126/sciadv.aba1306

5. Schulz TJ, Zarse K, Voigt A, Urban N, Birringer M, Ristow M. Glucose restriction extends Caenorhabditis elegans life span by inducing mitochondrial respiration and increasing oxidative stress. Cell Metabolism. 2007;6(4):280–293. doi:10.1016/j.cmet.2007.08.011

6. McCarter R, Mejia W, Ikeno Y, Monnier V, Kewitt K, Gibbs M, McMahan A, Strong R. Plasma glucose and the action of calorie restriction on aging. The Journals of Gerontology. Series A, Biological Sciences and Medical Sciences. 2007;62(10):1059–1070. doi:10.1093/gerona/62.10.1059

7. Galenza A, Hutchinson J, Campbell SD, Hazes B, Foley E. Glucose modulates Drosophila longevity and immunity independent of the microbiota. Biology Open. 2016;5(2):165–173. doi:10.1242/bio.015016

8. Garalde DR, Snell EA, Jachimowicz D, Sipos B, Lloyd JH, Bruce M, Pantic N, Admassu T, James P, Warland A, et al. Highly parallel direct RNA sequencing on an array of nanopores. Nature Methods. 2018;15(3):201–206. doi:10.1038/nmeth.4577

9. Montañés JC, Huertas M, Moro SG, Blevins WR, Carmona M, Ayté J, Hidalgo E, Albà MM. Native RNA sequencing in fission yeast reveals frequent alternative splicing isoforms. Genome Research. 2022;32(6):1215–1227. doi:10.1101/gr.276516.121

10. Thodberg M, Thieffry A, Bornholdt J, Boyd M, Holmberg C, Azad A, Workman CT, Chen Y, Ekwall K, Nielsen O, et al. Comprehensive profiling of the fission yeast transcription start site activity during stress and media response. Nucleic Acids Research. 2019;47(4):1671–1691. doi:10.1093/nar/gky1227

11. Ronne H. Glucose repression in fungi. Trends in Genetics. 1995;11(1):12–17. doi:10.1016/S0168-9525(00)88980-5

12. Hirota K, Miyoshi T, Kugou K, Hoffman CS, Shibata T, Ohta K. Stepwise chromatin remodelling by a cascade of transcription initiation of non-coding RNAs. Nature. 2008;456(7218):130–134.

13. Galipon J, Miki A, Oda A, Inada T, Ohta K. Stress-induced lncRNAs evade nuclear degradation and enter the translational machinery. Genes to Cells. 2013;18(5):353–368. doi:10.1111/gtc.12042

14. Oda A, Takemata N, Hirata Y, Miyoshi T, Suzuki Y, Sugano S, Ohta K. Dynamic transition of transcription and chromatin landscape during fission yeast adaptation to glucose starvation. Genes to Cells. 2015;20(5):392–407. doi:https://doi.org/10.1111/gtc.12229

15. Miki A, Galipon J, Sawai S, Inada T, Ohta K. RNA decay systems enhance reciprocal switching of sense and antisense transcripts in response to glucose starvation. Genes to Cells. 2016;21(12):1276–1289. doi:https://doi.org/10.1111/gtc.12443

16. Jia L, Mao Y, Ji Q, Dersh D, Yewdell JW, Qian S-B. Decoding mRNA translatability and stability from the 5′ UTR. Nature Structural & Molecular Biology. 2020;27(9):814–821. doi:10.1038/s41594-020-0465-x

17. Siegel DA, Le Tonqueze O, Biton A, Zaitlen N, Erle DJ. Massively parallel analysis of human 3′ UTRs reveals that AU-rich element length and registration predict mRNA destabilization. G3 Genes|Genomes|Genetics. 2022;12(1):jkab404. doi:10.1093/g3journal/jkab404

18. Toyoda Y, Saitoh S. Fission Yeast TORC2 Signaling Pathway Ensures Cell Proliferation under Glucose-Limited, Nitrogen-Replete Conditions. Biomolecules. 2021;11(10):1465. doi:10.3390/biom11101465

19. Welton RM, Hoffman CS. Glucose Monitoring in Fission Yeast via the gpa2 Gα, the git5 Gβ and the git3 Putative Glucose Receptor. Genetics. 2000;156(2):513–521. doi:10.1093/genetics/156.2.513

20. Takemata N, Oda A, Yamada T, Galipon J, Miyoshi T, Suzuki Y, Sugano S, Hoffman CS, Hirota K, Ohta K. Local potentiation of stress-responsive genes by upstream noncoding transcription. Nucleic Acids Research. 2016;44(11):5174– 5189. doi:10.1093/nar/gkw142

21. Soneson C, Love MI, Robinson MD. Differential analyses for RNA-seq: transcript-level estimates improve gene-level inferences. F1000Research. 2016;4:1521. doi:10.12688/f1000research.7563.2

22. Wood V, Harris MA, McDowall MD, Rutherford K, Vaughan BW, Staines DM, Aslett M, Lock A, Bähler J, Kersey PJ, et al. PomBase: a comprehensive online resource for fission yeast. Nucleic Acids Research. 2012;40(Database issue):D695–D699. doi:10.1093/nar/gkr853

23. Araujo PR, Yoon K, Ko D, Smith AD, Qiao M, Suresh U, Burns SC, Penalva LOF. Before It Gets Started: Regulating Translation at the 5′ UTR. International Journal of Genomics. 2012;2012:e475731. doi:10.1155/2012/475731

24. Kuersten S, Goodwin EB. The power of the 3′ UTR: translational control and development. Nature Reviews Genetics. 2003;4(8):626–637. doi:10.1038/nrg1125

25. Wu Q, Wright M, Gogol MM, Bradford WD, Zhang N, Bazzini AA. Translation of small downstream ORFs enhances translation of canonical main open reading frames. The EMBO Journal. 2020;39(17):e104763. doi:10.15252/embj.2020104763

26. Barbosa C, Peixeiro I, Romão L. Gene Expression Regulation by Upstream Open Reading Frames and Human Disease. PLoS Genetics. 2013;9(8):e1003529. doi:10.1371/journal.pgen.1003529

27. Cvijović M, Dalevi D, Bilsland E, Kemp GJ, Sunnerhagen P. Identification of putative regulatory upstream ORFs in the yeast genome using heuristics and evolutionary conservation. BMC Bioinformatics. 2007;8(1):295. doi:10.1186/1471-2105-8-295

28. Parker MT, Knop K, Sherwood AV, Schurch NJ, Mackinnon K, Gould PD, Hall AJ, Barton GJ, Simpson GG. Nanopore direct RNA sequencing maps the complexity of Arabidopsis mRNA processing and m6A modification Wan Y, Hardtke CS, editors. eLife. 2020;9:e49658. doi:10.7554/eLife.49658

29. Workman RE, Tang AD, Tang PS, Jain M, Tyson JR, Razaghi R, Zuzarte PC, Gilpatrick T, Payne A, Quick J, et al. Nanopore native RNA sequencing of a human poly(A) transcriptome. Nature Methods. 2019;16(12):1297–1305. doi:10.1038/s41592-019-0617-2

30. Takahashi H, Kato S, Murata M, Carninci P. CAGE-Cap Analysis Gene Expression: a protocol for the detection of promoter and transcriptional networks. Methods in molecular biology (Clifton, N.J.). 2012;786:181–200. doi:10.1007/978-1-61779-292-2_11

31. Li H. Aligning sequence reads, clone sequences and assembly contigs with BWA-MEM. 2013 [accessed 2023 Jul 1]. http://arxiv.org/abs/1303.3997. doi:10.48550/arXiv.1303.3997

32. Kim D, Paggi JM, Park C, Bennett C, Salzberg SL. Graph-based genome alignment and genotyping with HISAT2 and HISAT-genotype. Nature Biotechnology. 2019;37(8):907–915. doi:10.1038/s41587-019-0201-4

33. Leger A, Leonardi T. pycoQC, interactive quality control for Oxford Nanopore Sequencing. Journal of Open Source Software. 2019;4(34):1236. doi:10.21105/joss.01236

34. Li H. Minimap2: pairwise alignment for nucleotide sequences. Bioinformatics. 2018;34(18):3094–3100. doi:10.1093/bioinformatics/bty191

35. Danecek P, Bonfield JK, Liddle J, Marshall J, Ohan V, Pollard MO, Whitwham A, Keane T, McCarthy SA, Davies RM, et al. Twelve years of SAMtools and BCFtools. GigaScience. 2021;10(2):giab008. doi:10.1093/gigascience/giab008

36. Dobin A, Davis CA, Schlesinger F, Drenkow J, Zaleski C, Jha S, Batut P, Chaisson M, Gingeras TR. STAR: ultrafast universal RNA-seq aligner. Bioinformatics (Oxford, England). 2013;29(1):15–21. doi:10.1093/bioinformatics/bts635

37. Quinlan AR, Hall IM. BEDTools: a flexible suite of utilities for comparing genomic features. Bioinformatics. 2010;26(6):841–842. doi:10.1093/bioinformatics/btq033

38. Thodberg M, Thieffry A, Vitting-Seerup K, Andersson R, Sandelin A. CAGEfightR: analysis of 5′-end data using R/Bioconductor. BMC Bioinformatics. 2019;20(1):487. doi:10.1186/s12859-019-3029-5

39. Tang AD, Soulette CM, van Baren MJ, Hart K, Hrabeta-Robinson E, Wu CJ, Brooks AN. Full-length transcript characterization of SF3B1 mutation in chronic lymphocytic leukemia reveals downregulation of retained introns. Nature Communications. 2020;11(1):1438. doi:10.1038/s41467-020-15171-6

40. Tardaguila M, de la Fuente L, Marti C, Pereira C, Pardo-Palacios FJ, del Risco H, Ferrell M, Mellado M, Macchietto M, Verheggen K, et al. SQANTI: extensive characterization of long-read transcript sequences for quality control in full-length transcriptome identification and quantification. Genome Research. 2018;28(3):396–411. doi:10.1101/gr.222976.117

41. de la Fuente L, Arzalluz-Luque Á, Tardáguila M, del Risco H, Martí C, Tarazona S, Salguero P, Scott R, Lerma A, Alastrue-Agudo A, et al. tappAS: a comprehensive computational framework for the analysis of the functional impact of differential splicing. Genome Biology. 2020;21(1):119. doi:10.1186/s13059-020-02028-w

42. Tardaguila M, de la Fuente L, Marti C, Pereira C, Pardo-Palacios FJ, del Risco H, Ferrell M, Mellado M, Macchietto M, Verheggen K, et al. Running SQANTI3 filter. GitHub. 2023 May 11 [accessed 2023 Jun 15]. https://github.com/ConesaLab/SQANTI3/wiki/Running-SQANTI3-filter

43. Patro R, Duggal G, Love MI, Irizarry RA, Kingsford C. Salmon provides fast and bias-aware quantification of transcript expression. Nature Methods. 2017;14(4):417–419. doi:10.1038/nmeth.4197

44. Gleeson J, Leger A, Prawer YDJ, Lane TA, Harrison PJ, Haerty W, Clark MB. Accurate expression quantification from nanopore direct RNA sequencing with NanoCount. Nucleic Acids Research. 2022;50(4):e19. doi:10.1093/nar/gkab1129

45. Love MI, Huber W, Anders S. Moderated estimation of fold change and dispersion for RNA-seq data with DESeq2. Genome Biology. 2014;15(12):550. doi:10.1186/s13059-014-0550-8

46. Valero-Mora PM. ggplot2: Elegant Graphics for Data Analysis. Journal of Statistical Software. 2010;35:1–3. doi:10.18637/jss.v035.b01

47. Wu T, Hu E, Xu S, Chen M, Guo P, Dai Z, Feng T, Zhou L, Tang W, Zhan L, et al. clusterProfiler 4.0: A universal enrichment tool for interpreting omics data. The Innovation. 2021;2(3):100141. doi:10.1016/j.xinn.2021.100141

48. Yu G, Wang L-G, Yan G-R, He Q-Y. DOSE: an R/Bioconductor package for disease ontology semantic and enrichment analysis. Bioinformatics. 2015;31(4):608–609. doi:10.1093/bioinformatics/btu684

49. Ge SX, Jung D, Yao R. ShinyGO: a graphical gene-set enrichment tool for animals and plants. Bioinformatics. 2020;36(8):2628–2629. doi:10.1093/bioinformatics/btz931

50. Kang Y-J, Yang D-C, Kong L, Hou M, Meng Y-Q, Wei L, Gao G. CPC2: a fast and accurate coding potential calculator based on sequence intrinsic features. Nucleic Acids Research. 2017;45(W1):W12–W16. doi:10.1093/nar/gkx428

51. Mészáros B, Erdős G, Dosztányi Z. IUPred2A: context-dependent prediction of protein disorder as a function of redox state and protein binding. Nucleic Acids Research. 2018;46(W1):W329–W337. doi:10.1093/nar/gky384

52. Potter SC, Luciani A, Eddy SR, Park Y, Lopez R, Finn RD. HMMER web server: 2018 update. Nucleic Acids Research. 2018;46(W1):W200–W204. doi:10.1093/nar/gky448

53. Almagro Armenteros JJ, Tsirigos KD, Sønderby CK, Petersen TN, Winther O, Brunak S, von Heijne G, Nielsen H. SignalP 5.0 improves signal peptide predictions using deep neural networks. Nature Biotechnology. 2019;37(4):420–423. doi:10.1038/s41587-019-0036-z

54. Virtanen P, Gommers R, Oliphant TE, Haberland M, Reddy T, Cournapeau D, Burovski E, Peterson P, Weckesser W, Bright J, et al. SciPy 1.0: fundamental algorithms for scientific computing in Python. Nature Methods. 2020;17(3):261–272. doi:10.1038/s41592-019-0686-2

55. Dikbaş F. Compositional correlation analysis of gene expression time series. Academic Platform Journal of Engineering and Smart Systems. 2022;10(1):30–41. doi:10.21541/apjess.1060765

56. Kong AT, Leprevost FV, Avtonomov DM, Mellacheruvu D, Nesvizhskii AI. MSFragger: ultrafast and comprehensive peptide identification in mass spectrometry–based proteomics. Nature Methods. 2017;14(5):513–520. doi:10.1038/nmeth.4256

57. da Veiga Leprevost F, Haynes SE, Avtonomov DM, Chang H-Y, Shanmugam AK, Mellacheruvu D, Kong AT, Nesvizhskii AI. Philosopher: a versatile toolkit for shotgun proteomics data analysis. Nature Methods. 2020;17(9):869–870. doi:10.1038/s41592-020-0912-y

58. Yu F, Haynes SE, Nesvizhskii AI. IonQuant Enables Accurate and Sensitive Label-Free Quantification With FDR-Controlled Match-Between-Runs. Molecular & Cellular Proteomics. 2021;20:100077. doi:10.1016/j.mcpro.2021.100077

59. Demichev V, Szyrwiel L, Yu F, Teo GC, Rosenberger G, Niewienda A, Ludwig D, Decker J, Kaspar-Schoenefeld S, Lilley KS, et al. dia-PASEF data analysis using FragPipe and DIA-NN for deep proteomics of low sample amounts. Nature Communications. 2022;13(1):3944. doi:10.1038/s41467-022-31492-0

60. The UniProt Consortium. UniProt: the universal protein knowledgebase in 2021. Nucleic Acids Research. 2021;49(D1):D480–D489. doi:10.1093/nar/gkaa1100

61. Koopmans F, Li KW, Klaassen RV, Smit AB. MS-DAP Platform for Downstream Data Analysis of Label-Free Proteomics Uncovers Optimal Workflows in Benchmark Data Sets and Increased Sensitivity in Analysis of Alzheimer’s Biomarker Data. Journal of Proteome Research. 2023;22(2):374–386. doi:10.1021/acs.jproteome.2c00513

62. Pluskal T, Hayashi T, Saitoh S, Fujisawa A, Yanagida M. Specific biomarkers for stochastic division patterns and starvation-induced quiescence under limited glucose levels in fission yeast. The Febs Journal. 2011;278(8):1299–1315. doi:10.1111/j.1742-4658.2011.08050.x

63. Nguyen LAC, Inada T, Galipon J. Nanopore Direct RNA Sequencing of Monosome- and Polysome-Bound RNA. In: Arakawa K, editor. Nanopore Sequencing: Methods and Protocols. New York, NY: Springer US; 2023. p. 281–297. (Methods in Molecular Biology). https://doi.org/10.1007/978-1-0716-2996-3_20. doi:10.1007/978-1-0716-2996-3_20

64. Magistri M, Faghihi MA, St. Laurent G, Wahlestedt C. Regulation of Chromatin Structure by Long Noncoding RNAs: Focus on Natural Antisense Transcripts. Trends in genetics : TIG. 2012;28(8):389–396. doi:10.1016/j.tig.2012.03.013

65. Brown T, Howe FS, Murray SC, Wouters M, Lorenz P, Seward E, Rata S, Angel A, Mellor J. Antisense transcription-dependent chromatin signature modulates sense transcript dynamics. Molecular Systems Biology. 2018;14(2):e8007. doi:10.15252/msb.20178007

66. Muskovic W, Slavich E, Maslen B, Kaczorowski DC, Cursons J, Crampin E, Kavallaris M. High temporal resolution RNA-seq time course data reveals widespread synchronous activation between mammalian lncRNAs and neighbouring protein-coding genes. Genome Research. 2022 Jun 27:gr.276818.122. doi:10.1101/gr.276818.122

67. Wery M, Gautier C, Descrimes M, Yoda M, Vennin-Rendos H, Migeot V, Gautheret D, Hermand D, Morillon A. Native elongating transcript sequencing reveals global anti-correlation between sense and antisense nascent transcription in fission yeast. RNA. 2018;24(2):196–208. doi:10.1261/rna.063446.117

68. Clément-Ziza M, Marsellach FX, Codlin S, Papadakis MA, Reinhardt S, Rodríguez-López M, Martin S, Marguerat S, Schmidt A, Lee E, et al. Natural genetic variation impacts expression levels of coding, non-coding, and antisense transcripts in fission yeast. Molecular Systems Biology. 2014;10(11):764. doi:10.15252/msb.20145123

69. Leong HS, Dawson K, Wirth C, Li Y, Connolly Y, Smith DL, Wilkinson CRM, Miller CJ. A global non-coding RNA system modulates fission yeast protein levels in response to stress. Nature Communications. 2014;5(1):3947. doi:10.1038/ncomms4947

70. Zheng D, Wang R, Ding Q, Wang T, Xie B, Wei L, Zhong Z, Tian B. Cellular stress alters 3′UTR landscape through alternative polyadenylation and isoform-specific degradation. Nature Communications. 2018;9(1):2268. doi:10.1038/s41467-018-04730-7

71. Urso SJ, Sathaseevan A, Brent Derry W, Lamitina T. Regulation of the hypertonic stress response by the 3′ mRNA cleavage and polyadenylation complex. Genetics. 2023;224(1):iyad051. doi:10.1093/genetics/iyad051

72. Wang T, Ye W, Zhang J, Li H, Zeng W, Zhu S, Ji G, Wu X, Ma L. Alternative 3′-untranslated regions regulate high-salt tolerance of Spartina alterniflora. Plant Physiology. 2023;191(4):2570–2587. doi:10.1093/plphys/kiad030

73. Woo YM, Kwak Y, Namkoong S, Kristjánsdóttir K, Lee SH, Lee JH, Kwak H. TED-Seq Identifies the Dynamics of Poly(A) Length during ER Stress. Cell Reports. 2018;24(13):3630–3641.e7. doi:10.1016/j.celrep.2018.08.084

74. Wu X, Wang J, Wu X, Hong Y, Li QQ. Heat Shock Responsive Gene Expression Modulated by mRNA Poly(A) Tail Length. Frontiers in Plant Science. 2020 [accessed 2022 Nov 8];11. https://www.frontiersin.org/articles/10.3389/fpls.2020.01255

75. Yan C, Wang Y, Lyu T, Hu Z, Ye N, Liu W, Li J, Yao X, Yin H. Alternative Polyadenylation in response to temperature stress contributes to gene regulation in Populus trichocarpa. BMC Genomics. 2021;22(1):53. doi:10.1186/s12864-020-07353-9

76. Mayr C, Bartel DP. Widespread shortening of 3’UTRs by alternative cleavage and polyadenylation activates oncogenes in cancer cells. Cell. 2009;138(4):673–684. doi:10.1016/j.cell.2009.06.016

77. Navarro E, Mallén A, Hueso M. Dynamic Variations of 3′UTR Length Reprogram the mRNA Regulatory Landscape. Biomedicines. 2021;9(11):1560. doi:10.3390/biomedicines9111560

78. Hong D, Jeong S. 3’UTR Diversity: Expanding Repertoire of RNA Alterations in Human mRNAs. Molecules and Cells. 2023;46(1):48–56. doi:10.14348/molcells.2023.0003

79. Callahan KP, Butler JS. TRAMP Complex Enhances RNA Degradation by the Nuclear Exosome Component Rrp62. Journal of Biological Chemistry. 2010;285(6):3540–3547. doi:10.1074/jbc.M109.058396

80. Eckmann CR, Rammelt C, Wahle E. Control of poly(A) tail length. Wiley Interdisciplinary Reviews: RNA. 2011;2(3):348–361. doi:10.1002/wrna.56

81. Liu Y, Zhao H, Shao F, Zhang Y, Nie H, Zhang J, Li C, Hou Z, Chen Z-J, Wang J, et al. Remodeling of maternal mRNA through poly(A) tail orchestrates human oocyte-to-embryo transition. Nature Structural & Molecular Biology. 2023;30(2):200–215. doi:10.1038/s41594-022-00908-2

82. Xiang K, Bartel DP. The molecular basis of coupling between poly(A)-tail length and translational efficiency Nilsen TW, Manley JL, Nilsen TW, editors. eLife. 2021;10:e66493. doi:10.7554/eLife.66493

83. Zheng F, Jia B, Dong F, Liu L, Rasul F, He J, Fu C. Glucose starvation induces mitochondrial fragmentation depending on the dynamin GTPase Dnm1/Drp1 in fission yeast. Journal of Biological Chemistry. 2019;294(47):17725–17734. doi:10.1074/jbc.RA119.010185

84. Buljan M, Bateman A. The evolution of protein domain families. Biochemical Society Transactions. 2009;37(4):751–755. doi:10.1042/BST0370751

85. Nasir A, Kim KM, Caetano-Anollés G. Global Patterns of Protein Domain Gain and Loss in Superkingdoms. PLOS Computational Biology. 2014;10(1):e1003452. doi:10.1371/journal.pcbi.1003452

86. Kress A, Poch O, Lecompte O, Thompson JD. Real or fake? Measuring the impact of protein annotation errors on estimates of domain gain and loss events. Frontiers in Bioinformatics. 2023 [accessed 2023 Jul 1];3. https://www.frontiersin.org/articles/10.3389/fbinf.2023.1178926

87. Beyer A, Hollunder J, Nasheuer H-P, Wilhelm T. Post-transcriptional expression regulation in the yeast Saccharomyces cerevisiae on a genomic scale. Molecular & cellular proteomics: MCP. 2004;3(11):1083–1092. doi:10.1074/mcp.M400099-MCP200

88. Koussounadis A, Langdon SP, Um IH, Harrison DJ, Smith VA. Relationship between differentially expressed mRNA and mRNA-protein correlations in a xenograft model system. Scientific Reports. 2015;5(1):10775. doi:10.1038/srep10775

89. Nie L, Wu G, Zhang W. Correlation between mRNA and protein abundance in Desulfovibrio vulgaris: A multiple regression to identify sources of variations. Biochemical and Biophysical Research Communications. 2006;339(2):603–610. doi:10.1016/j.bbrc.2005.11.055

90. Upadhya SR, Ryan CJ. Experimental reproducibility limits the correlation between mRNA and protein abundances in tumor proteomic profiles. Cell Reports Methods. 2022;2(9):100288. doi:10.1016/j.crmeth.2022.100288

91. Gatti L, Hoe KL, Hayles J, Righetti SC, Carenini N, Bo LD, Kim DU, Park HO, Perego P. Ubiquitin-proteasome genes as targets for modulation of cisplatin sensitivity in fission yeast. BMC Genomics. 2011;12(1):44. doi:10.1186/1471-2164-12-44

