## Supplementary Tables description for "Functional consequences of shifting transcript boundaries in glucose starvation"

Supplementary Table 1: All the genes obtained with FLAIR/SQANTI3 pipeline after SQANTI3 quality filtering with the number of isoforms associated to each gene

Supplementary Table 2: Description of the genes being in both SQANTI and Pombase, SQANTI or Pombase only

Supplementary Table 3: Description of the DEGs by cluster of their gene expression

Supplementary Table 4: Description of the DEGs by category as described in Figure 3d

Supplementary Table 5: Mean of the gene expression for each gene at each time point into glucose starvation with their associated antisense’s and the correlation score in both non-lag and lag scenarios

Supplementary Table 6: Description of the genes being both a DEG and a DTU, a DEG or a DTU only

Supplementary Table 7: Description of the DEGs by category as described in Figure 4b

Supplementary Table 8: Description of the genes that have a change in poly(A) tail length during glucose starvation

Supplementary Table 9: Description of the DTUs with domain gains and losses with the domains affected

Supplementary Table 10: Description of the DTUs with domain gains and losses with the coverage of the domains normalized by the total protein coverage to observe whether a domain gain/loss as defined in Supplementary Table 9 is actually happening or not

Supplementary Table 11: Description of the DDPs by cluster of their protein abundance

Supplementary Table 12: Description of the DDPs by category as described in Figure 5c

Supplementary Table 13: Description of the DEGs, DTUs and DDPs and the different overlaps
